# Caspase-Activated DNase localizes to cancer causing translocation breakpoints during cell differentiation

**DOI:** 10.1101/2024.09.24.614809

**Authors:** Dalal Alsowaida, Brian D. Larsen, Sarah Hachmer, Mehri Azimi, Eric Arezza, Steve Brunette, Steven Tur, Carmen G. Palii, Bassam Albraidy, Claus S. Sorensen, Marjorie Brand, F. Jeffrey Dilworth, Lynn A. Megeney

## Abstract

Caspase activated DNase (CAD) induced DNA breaks promote cell differentiation and therapy-induced cancer cell resistance. CAD targeting activity is assumed to be unique to each condition, as differentiation and cancer genesis are divergent cell fates. Here, we made the surprising discovery that a subset of CAD-bound targets in differentiating muscle cells are the same genes involved in the genesis of cancer-causing translocations. In muscle cells, a prominent CAD-bound gene pair is *Pax7* and *Foxo1a*, the mismatched reciprocal loci that give rise to alveolar rhabdomyosarcoma. We show that CAD-targeted breaks in the *Pax7* gene are physiologic to reduce *Pax7* expression, a prerequisite for muscle cell differentiation. A cohort of these CAD gene targets are also conserved in early differentiating T cells and include genes that spur leukemia/lymphoma translocations. Our results suggest the CAD targeting of translocation prone oncogenic genes is non-pathologic biology and aligns with initiation of cell fate transitions.

## Main

Caspase activated DNase (CAD) is the terminal enzyme in apoptosis that initiates dissolution of chromatin by inflicting genome wide DNA double strand breaks^1^. CAD activity is normally constrained by physical interaction with a nascent protein inhibitor ICAD (Inhibitor of Caspase Activated DNase), which is released from CAD by a direct caspase 3 cleavage event when apoptosis is engaged^1^. In contrast to the unconstrained activation of CAD during cell death, limited activation of this nuclease, through transient activation of caspase 3, has been implicated in promoting nonlethal cell fate decisions^2^. For example, differentiation of murine skeletal muscle progenitor cells and Drosophila macrophage progenitor cells are both dependent on CAD mediated DNA strand breaks^3,4^. CAD genome targets have yet to be identified in the Drosophila macrophage lineage^4^, however, in muscle progenitor cells, CAD mediated breaks have been mapped to the p21 gene promoter, a critical cell cycle regulatory factor. In turn, this CAD directed DNA damage event primes p21 gene expression, which is essential for effective initiation of the differentiation program^3^.

In addition to cell fate instruction, reports suggest that CAD activity enhances cancer cell fitness and survival. Studies have reported that sub-lethal activation of caspase 3, through engagement of the mitochondrial cell death pathway, is synonymous with cancer cell growth and metastasis^5,6,7,8^. This protease dependency for cancer cell adaptation has been directly linked to caspase 3 targeted cleavage of ICAD and release of CAD, which appears to induce genome wide DNA damage and micro-nuclei formation^9,10^. An obvious interpretation of these studies is that these DNA strand breaks are oncogenic, yet precise genetic targets of CAD that sustain and amplify cell transformation have not been identified. CAD mediated DNA strand breaks also compel the emergence of radiotherapy induced resistance in cancer cells/tumors. Here, the mechanism at play is unique, caspase 3 independent, and originates from direct phosphorylation of ICAD, which then acts to physically recruit CAD to the genome, activating its nuclease function rather than repressing it^11^. The resulting CAD induced DNA strand breaks target CTCF enriched binding sites that promote cell cycle withdrawal, allowing the cancer cells to exit the cell cycle, repair the DNA damage, avoiding the radiation sensitivity that characterizes actively dividing cancer cells^11^.

These observations support the contention that CAD is a potent genome modifier with the capacity to alter cell fate independent of cell death. The obvious subtext to this hypothesis is that CAD must target unique genome regions to achieve divergent cell fates, such as differentiation versus cancer cell maintenance and expansion. Nevertheless, CAD may target similar genomic loci irrespective of cell fate, where the precise cellular context determines whether the strand breaks in question activate or repress gene expression. To clarify and refine our understanding of CAD as a cell fate reprogramming nuclease, we performed CUT&Tag against endogenous CAD in differentiating skeletal muscle cells. CAD displayed a high degree of binding to promoters and enhancers of genes which change significantly during cell differentiation, a cohort of which are robustly elevated. We also identified a subset of CAD bound targets in differentiating muscle cells and early differentiating T cells to be the same genes involved in the genesis of cancer-causing breakpoint translocations. The CAD targeting of breakpoint prone genes is over-represented by genes that must be repressed for the differentiation program to proceed. Together these results suggest that CAD activity which compels normal cell differentiation may also produce strand breaks in parent genes that are susceptible to the eventual formation of oncogenic breakpoint translocations.

## Results

### CAD binds gene promoters and is associated with elevated gene expression in muscle cell differentiation

Prior investigations of CAD DNA binding in cancer cells indicated low abundance genome interactions^11^ suggesting a probable divergent targeting activity compared to the more robust degree of CAD inflicted DNA strand breaks we observed during muscle cell differentiation^3,12^. Therefore to assess CAD genome wide targeting preferences in differentiating C2C12 muscle cells, and to further our understanding of the cell fate modifying activity for this nuclease, we performed cleavage under targets and tagmentation (CUT&Tag) against endogenous CAD protein on DNA. To improve CAD binding retention on ambient DNA targets, C2C12 cells were also treated with a caspase 3 peptide inhibitor. The rationale for this adjustment is that prior experiments indicated CAD and its protein inhibitor ICAD are constitutively bound to DNA, becoming more labile once CAD is activated through caspase 3 cleavage of ICAD^13^. Accordingly, CUT&Tag profiling revealed robust CAD genome binding in caspase 3 inhibited C2C12 muscle cells compared to cells with intact caspase 3 activity (peak height/binding intensity in growth vs 24 hours differentiation) as mapped using computeMatrix function in deepTools^14^ (Fig. 1a; 1b), although the total number of binding sites did not vary as determined by the average peak call across the data using MACS, SERCS and GoPeaks. Genome distribution of the binding sites revealed a concentration of CAD binding at or near transcription start sites (TSS) (Fig. 1a; 1b), with binding site preferences for repetitive regions (Fig. 1c; Extended Data Fig. 1a). Use of MACS and the annotatePeak and getPromoters function in the ChIPseeker program revealed marked gene specific distribution for CAD, with 43.32% of the total CAD binding sites found within core promoters (<1 KB from the TSS) (Fig. 1d). This promoter based CAD binding preference was reminiscent of prior observations from our group, where we noted single strand breaks in the p21 gene that were synonymous with increased p21 gene expression^3^.

**Figure 1.**
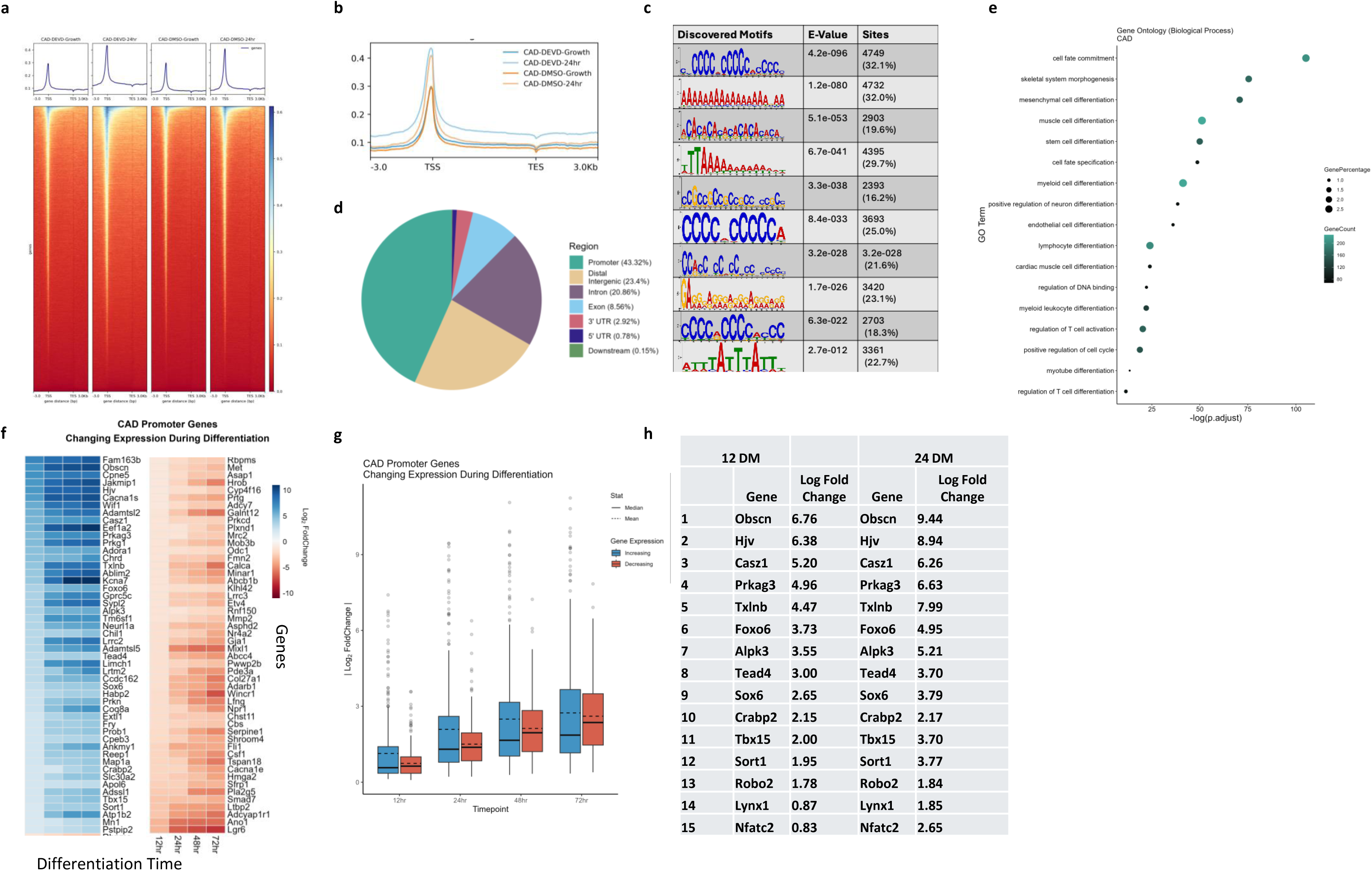
Caspase Activated DNase (CAD) displays a binding preference for promoters of differentiation expressed genes. <u>Genomic distribution of CAD signal for CAD binding sites.</u> (A) plotHeatmaps of CUT and Tag show count per million (CPM) normalized signals for reads across 3,000 bp genomic intervals relative to the TSS. Genomic distribution of the binding sites of CAD at Growth, and 24 hours post differentiation from C2C12 muscle cells treated with DMSO or z.DEVD-FMK. Data sets were generated from merged, two independent biological replicates for each condition. Heatmap profiles were generated using deepTools computeMatrix^52^. Peak calling of binding sites was performed using Model-based Analysis of Chip-Seq. (MACS), Sparse Enrichment Analysis for CUT & RUN (SEACR), and GoPeaks. (B) PlotProfile averaged profiles of the compressed matrix of occupancy/signal near TSS produced by computeMatrix. (C) Graphical representation of sequence motifs detected in CAD DNA sequences using Motif enrichment analysis, demonstrates repetitive element enrichment for CAD bound genomic regions (as calculated with STREME; E-values of motifs are indicated). (D) Pie chart illustrating percent genomic distribution of the binding sites of CAD at 24 hours post differentiation from C2C12 muscle cells treated with z.DEVD-FMK, indicates enrichment at promoters, enhancers, distal intergenic regions, exons, introns, two untranslated regions (UTRs). The DiffBind tool was used to obtain the consensus peakset and AnnotatePeak function was used with the ChIPseeker package to determine peaks. (E) Gene Ontology (GO) analysis identified within CAD bound loci for biological processes demonstrates enrichment of genes that regulate gene expression and chromatin modification. <u>RNA-Seq. analysis of CAD promoter genes.</u> (F) Heatmaps depicting changes in gene expression for CAD promoter bound gene targets during 12-, 24-, 48-, and 72-hours differentiation relative to growth conditions (reference point). Blue color indicates elevated mRNA/gene expression, while red color indicates decreased mRNA/gene expression levels (Color scale represents the log2fold change). Three biological replicates for each timepoint were assessed, and DESeq2 results were obtained to get the log2FoldChange and p values. Genes with p. adjusted < 0.05 across all timepoint comparisons were considered significant for expression trend analysis. (G) Box-plot diagram depicts the distributions of gene expression log2Fold change across different differentiation timepoints 12, 24, 48, and 72 hours. The central line in the box marking the median value. (H) The subset of CAD bound genes (promoters) are listed with an increase in gene expression in C2C12 muscle cells at 12- and 24-hour differentiating cells (single gene data points from the box plots in G that are above the median) All genes were filtered first that the adjusted p-value for all timepoints is < 0.05. Then the median was taken for each timepoint 12 and 24 hours to be equal to or greater than 2 log2fold value.

GO and KEGG pathway analysis of the CAD bound genome indicated that bound genes were enriched for functions related to regulation of multi-lineage cell differentiation, gene expression, etc (Fig. 1e; Extended Data Fig. 1b), consistent with a role for CAD in modifying cell fate decisions. In addition, peak calling across these individual promoters combined with RNAseq analysis of differentiating C2C12 muscle cells revealed that CAD bound promoters display a broadly similar number of decreased and increased expressed genes (Fig. 1f; Extended Data Fig. 1c). The cohort of decreased expressed genes in differentiating muscle cells is likely reflective of CAD induced double strand breaks (DSBs), which are known to cause transcription repression^15^. However, the cohort of increased expressed genes bound by CAD displayed a large number of genes that were elevated significantly above the mean, implying a unique CAD gene induction activity (Fig. 1g). Analysis of these CAD bound super-responder genes indicated well known differentiation inducing factors, such as *Casz1* and *Tead4* (Fig. 1h). We also observed significant CAD binding within the p21 gene, which was distal to the single strand breaks in the p21 promoter we had observed in prior experiments. These strand breaks appear similarly inductive for p21 transcription^3^, with escalating p21 expression from 2 to 3.5 fold above growth conditions. As such CAD may act as an initiation cue for cell differentiation by inflicting genome wide single strand breaks on responsive promoters/regulatory regions, which then act in concert to reprogram cell fate.

### CAD binding is enriched on parent genes that comprise break point translocations (BPTs)

The CUT&Tag analysis also identified a common CAD binding event in differentiating muscle cells (29.4% of the total CAD peaks called), specifically in introns and exons (Fig. 1d). This concentration of CAD within gene bodies suggests a second function for the nuclease during cell differentiation, where DNA strand breaks within exons and introns are less likely to be gene inductive events and are more inclined to create structural impediments for gene expression, as noted above^15^. The Initial curation and inspection of the CAD intragenic targets also revealed enrichment of genes that give rise to oncogenic breakpoint translocations (BPTs), where breaks in individual genes are mismatched with other broken genes in the subsequent DNA repair process, forming novel fusion genes^16,17,18,19^. These CAD targets include a broad range of known oncogenic BPTs, including TLX genes, *Nkx2.1*, *LMO2*, etc. (Extended Data Fig. 2a). CAD targeting of BPT prone genes during cell differentiation may be limited in scope, impacting a small subset of genes, or CAD may range across the genome, targeting all BPT sensitive loci. To address the relative frequency of CAD targeting to BPT affected genes in differentiating murine muscle cells, we conducted a direct comparison of the CAD bound genome against the known compendium of described human cancer causing BPTs, as curated by the Catalogue Of Somatic Mutations In Cancer (COSMIC; https://cancer.sanger.ac.uk/cosmic/about). The degree of CAD association with the COSMIC human BPT genes was very high in differentiating mouse muscle cells, with CAD bound to ∼53.4% of the total BPT genes identified (183 out of 343 genes) (UpSet plot Fig. 2a; Circablot Fig. 2b). This cross species CAD gene binding in murine muscle cells matched to human BPTs was also characterized by a significant degree of precise syntenic site overlap between CAD and the gene breakpoints (Fig. 2b, 2c; 100 CAD bound sites directly overlap in 69 of the 183 CAD bound genes). Interestingly, CAD binding to the BPT genes had distinct motif binding preferences, which was dissimilar to the repetitive binding motifs found across intragenic and promoter regions (Extended Data Fig. 2c), suggesting that CAD BPT prone regions may share more defining characteristics, compared to other CAD bound regions.

**Figure 2.**
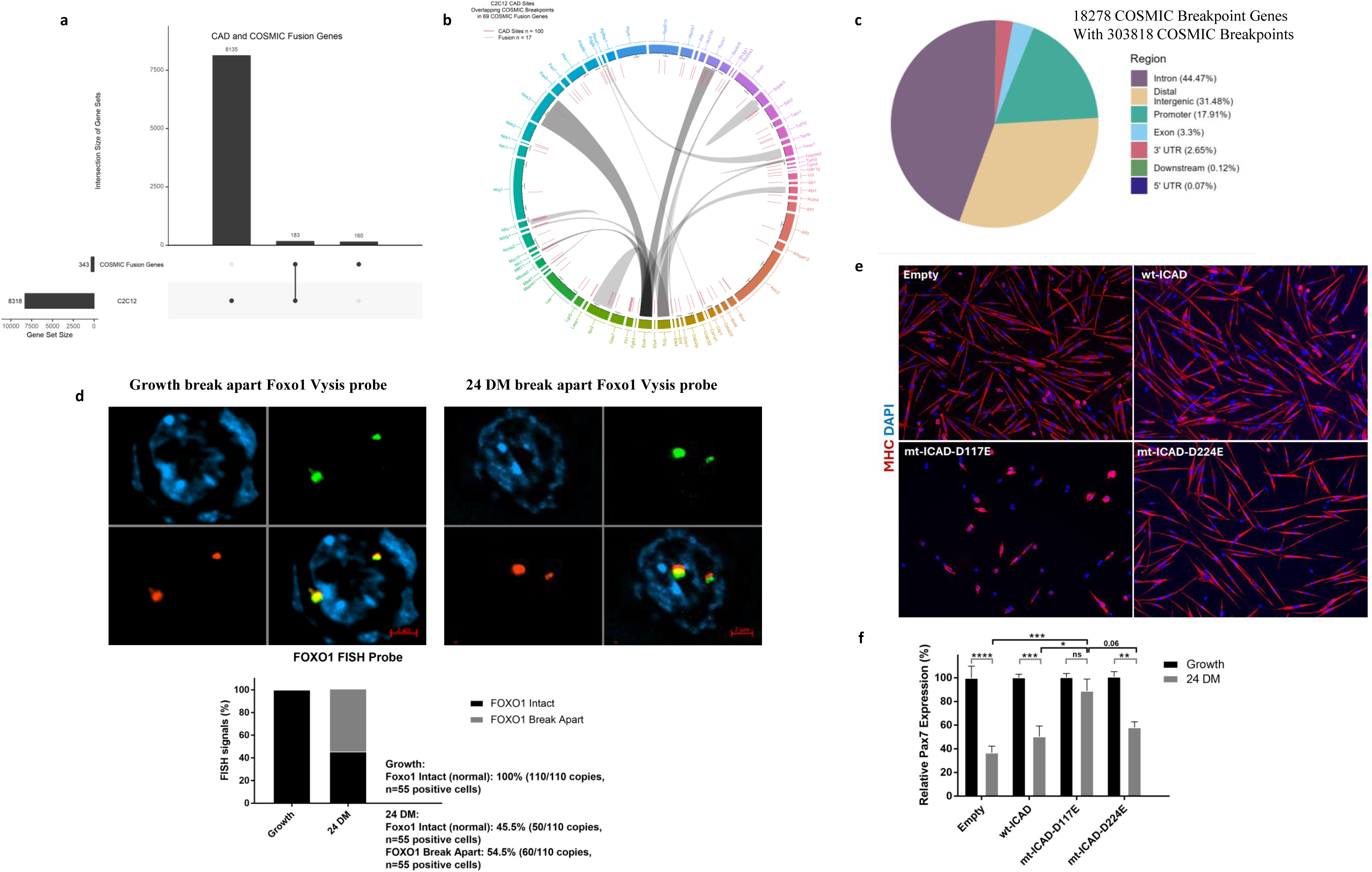
CAD is enriched at known cancer breakpoint prone genes and may target these loci to alter expression and propel cell differentiation. <u>CAD relationship to cancer causing breakpoint translocation genes.</u> (A) Upset plot visualizes the intersection size of gene sets between CAD bound loci from C2C12 muscle cells with breakpoint translocation/fusion genes from COSMIC database. COSMIC fusion genes were mapped to the mm10 orthologs from bioMart and obtained the intersection and union of the CAD and COSMIC genes to plot the Upset plot. CAD bound to ∼53.4% of the total BPT genes identified (183 out of 343 genes) (UpSet plot Fig. 2a). (B) Circa plot visualizing cross-comparison between CAD-binding sites and COSMIC fusion genes set. Tracks from inside to the outside correspond to (1) gene synteny between chromosomes, (2) genes For the Circa plot, from outer to inner: Rainbow: represents COSMIC fusion genes for which CAD sites are bound (i.e. CAD-Fusion genes), Red/teal/purple lines represents CAD site locations where COSMIC Breakpoints overlap with the CAD sites (i.e. CAD-Breakpoints) in the fusion genes, lined-up with their respective genes in their gene ranges (and coloured). CAD gene binding in murine muscle cells matched to human BPTs displayed a significant degree of precise syntenic site overlap between CAD and the gene breakpoints (Fig. 2b, 2c; 100 CAD bound sites directly overlap in 69 of the 183 CAD bound genes). The innermost links “Grey” indicate: Gene fusion pairs for CAD-Fusion genes in the circus. Circa plots were made with the circlize package (R package). (C) Pie chart representing genomic distribution of COSMIC breakpoint genes within breakpoints. <u>CAD targets and breaks Pax7 and Foxo1 loci to reduce expression of these genes and advance</u> <u>muscle cell differentiation.</u> (D) FISH analysis to Foxo1 gene integrity in primary myoblast cells during cell growth and 24 hours post differentiation. The FISH probe is designed to detect break apart gene loci involving the *FOXO1* (FKH1, FKHR) gene located on chromosome 13q14. For Foxo1 FISH probe, 110 copies (n=55) of intact FISH probes (100%) were quantified during growth conditions. In differentiated muscle cells (n=55), 50 copies of intact FISH probes (45.5%) and 60 copies of break apart FISH signals (54.5%) were quantified. (E) Primary myoblasts expressing wt-hICAD, mt-hICAD-D117E, and mt-hICAD D224E were subject to low serum induction of differentiation for 72 hours, which was monitored with staining for the differentiation specific marker, myosin heavy chain (MHC, red) nuclei were counterstained with DAPI (blue). (F) The same ICAD constructs were used to generate stable transfectants in primary myoblasts, where hICAD-D117E transfected cells (repressed CAD activity) maintained elevated Pax7 gene expression post low serum induction of differentiation. Relative expression of Pax7 at 24 hours following the induction of differentiation was determined and normalized to the Pax7 expression in each individual cell line at growth. n=4 (in triplicate). *: p<0.05, **: p<0.01, ***: p<0.001****: p<0.0001 using two-way ANOVA-Tukey’s multiple comparisons test.

### CAD targeting of the rhabdomyosarcoma BPT gene Pax7 is a required to promote normal muscle cell differentiation

These data suggest that the CAD nuclease is positioned within the genome to inflict breaks that preconfigure BPT formation. Similar to CAD bound promoters, CAD targeting of BPT sensitive genes may be part of inducing the differentiation program as noted above, i.e. breaking a gene through a DSB to reduce its expression. This is a reasonable assumption, as it is very unlikely that BPTs would be a conserved alteration solely dedicated to the induction of a cancer mutation per se. Cross comparison of CAD bound BPT genes against the RNAseq data revealed that a number of these CAD target genes have decreased levels of expression in differentiating muscle cells versus growing cells (Extended Data Fig. 2b). One CAD bound BPT pair of particular interest was *Pax7* and *Foxo1a*. Breaks in the *Pax7* and *Foxo1a* genes give rise to reciprocal translocations producing a fusion gene with robust oncogenic activity that forms aggressive solid tumours, referred to as alveolar rhabdomyosarcomas (ARMs)^20,21^. The oncogenic activity in the Pax7:Foxo1 fusion protein is hypothesized to arise from a self re-inforcing phenomenon, where the fusion proteins are resistant to normal degradation events that target Pax proteins in muscle stem/progenitor cells, propelling cell growth^22,23^. In a normal physiologic context, Pax7 confers a muscle stem/progenitor cell identity^24^, yet *Pax7* gene expression must decrease to allow differentiation to proceed^25,26^. We noted that CAD binding varied considerably at the *Pax7*/*Foxo1a* loci between growth vs differentiation, the latter being very prominent (Extended Data Fig. 2d). Moreover, CAD bound to the exon intron boundary in murine C2C12 muscle cells that is syntenic with the breakpoint in the human *Pax7* gene that propels aRMS. FISH analysis using probes to these regions determined that the Foxo1a gene was subject to breakage during normal primary muscle cell differentiation but not in growth conditions (Fig. 2d). In addition, the full translocation event between the *Pax7* and *Foxo1* loci could be detected by PCR/qPCR at the DNA and transcript level (respectively), as well as with the use of FISH to detect *Pax7*/*Foxo1a* fusion genes in differentiating C2C12 and primary derived skeletal muscle cells (Extended Data Fig. 2e; Extended Data Fig. 2f).

Next, we determined whether caspase 3 induced CAD activation was required to directly repress *Pax7* gene expression. To this end, expression of a caspase 3 cleavage mutant of ICAD (D117E) (which we have shown dramatically curtails CAD induced strand breaks)^3^, led to de-repression of *Pax7* expression and impaired muscle cell differentiation (Fig. 2e, 2f). These results suggest that CAD targets the gene breaks that configure the aRMS BPT, and that this targeting event is normal biology, where the CAD induced DSB break reduces *Pax7* gene expression to allow muscle cell differentiation to proceed. This observation implies that a differentiating progenitor cell may be the source of the initial DSBs that lead to BPTs.

### CAD targets promoters and BPT prone genes in differentiating T cells

We noted that a large cohort of the known leukemic and lymphomic BPT parent genes were bound by CAD in differentiating muscle cells (Extended Data Fig. 2a) suggesting that CAD may drive translocation formation beyond sarcomas. The maturation of this hematopoietic lineage is known to be dependent on caspase 3 activity^27,28^ and stem cells in this compartment are also considered to be the source of the BPT transformed cells that seed leukemias^29^. As such, we characterized CAD genome wide binding in early differentiating T cells. We used single positive and double negative selection of primary derived mouse thymic progenitors (Thy1.2^+^CD4^−^CD8^−^) to isolate early differentiating T cell progenitors (Fig. 3a), then used the same experimental conditions to perform CUT&Tag against endogenous immune-precipitated CAD protein. The CAD bound genome in early differentiating T cell progenitors displayed similar promoter/enhancer and gene body distribution as that noted for CAD in differentiating skeletal muscle cells (Fig. 3b vs Fig. 1d), which was similar to the gene ontology analysis for the CAD bound genome in both cell lineages, i.e. genes related to chromatin modification and control of gene expression (Fig. 1e). Cross comparison of the CAD bound genome against the human BPT atlas revealed that CAD bound to a significant cohort of the parent translocation genes also bound in C2C12 muscle cells, with a higher total CAD interaction to fusion genes in T cells (Fig. 3c; CAD bound to 261 of the 343 total fusion genes). However, the overlap in binding between human BPTs and CAD was similar in each differentiating cell type (see Circablot Fig 2b). For example, T cells displayed CAD binding to 59 of the 343 BPT genes in the COSMIC database, with 62 exact overlaps on breaks in 56 CAD bound fusion genes, suggesting a similar degree of active BPT targeting across unrelated cell progenitors lineages (Fig.3c; Extended Fig. 3a).

**Figure 3.**
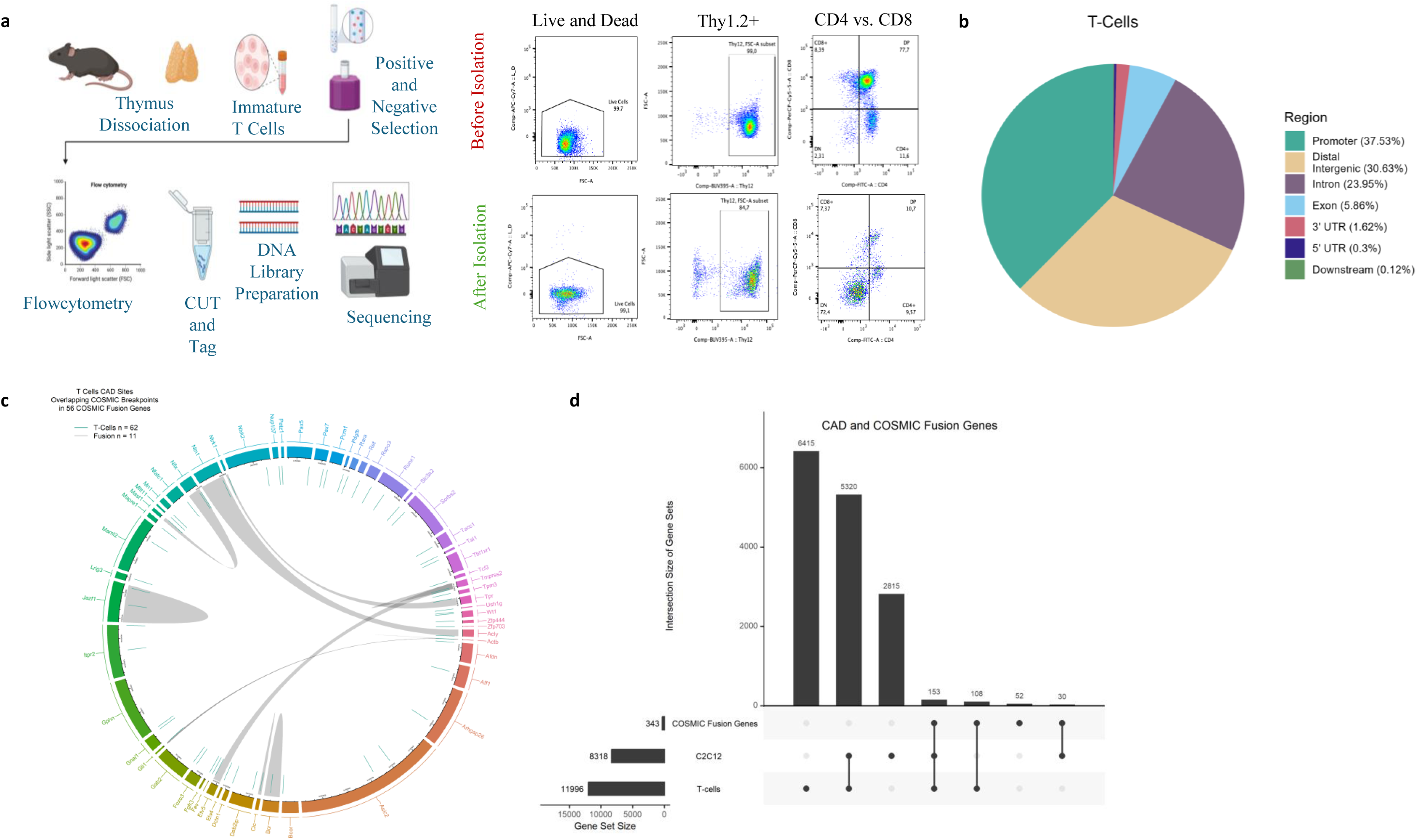
CAD binds to genomic sites in murine T cell progenitors that align with a subset of human breakpoint translocation genes. Schematic representation of the workflow for the CUT and Tag protocol applied to T cells isolated from thymus of wild type mice. Cell suspensions were acquired from mouse thymus then subject to FACS purification (Thy1.2^+^CD4^−^CD8^−^) to isolate early differentiating T cell progenitors (B) Pie chart illustrating the genomic distribution of CAD binding sites in differentiating T cell progenitors, showing the percentage for each genomic location category. Enrichment was noted at core promoters, distal intergenic regions, introns, exons, with a limited distribution at 3’ and 5’ untranslated regions (UTRs). The DiffBind tool was used to obtain the consensus peakset and AnnotatePeak function was used with the ChIPseeker package to determine peaks. (C) Circa plot visualizing the cross-comparison of CAD-binding sites between T cells and the COSMIC fusion gene set. Tracks from inside to the outside correspond to (1) gene synteny between chromosomes, (2) genes For the Circa plot, from outer to inner: Rainbow: represents COSMIC fusion genes for which CAD sites are bound (i.e. CAD-Fusion genes), Red/teal/purple lines represents CAD site locations where COSMIC Breakpoints overlap with the CAD sites (i.e. CAD-Breakpoints) in the fusion genes, lined-up with their respective genes in their gene ranges (and coloured). The innermost links “Grey” indicate: Gene fusion pairs for CAD-Fusion genes in the circus. Circa plots were made with the circulize package (R package). (D) UpSet plot visualizes the intersection size of gene sets between CAD from C2C12 muscle cells and fusion genes from T cell progenitors and the COSMIC fusion gene set.

## Discussion

Here, we establish that caspase activated DNase (CAD) targets both promoters and gene bodies to reorient gene expression in the promotion of cell differentiation. Our observations indicate that CAD targeting of promoters is generally a gene induction event, whereas CAD binding to gene bodies is consistent with a gene repression event. This latter activity is of particular relevance to cancer biology as CAD binding at gene bodies is enriched at break point translocation (BPT) prone genes. BPTs are a common mutation class in the etiology of human cancer, with curated examples that range from 300+ defined fusion loci that produce in-frame BPTs of different combinations to yield 800+ fusion transcripts, all having defined gain of function oncogenic properties (https://ccsm.uth.edu/FusionGDB/). The frequency and non-random recurrence of BPTs across the human population suggests that affected genome regions may be predisposed to damage/breakage. Accordingly, several hypothesis have emerged, implying BPT prone regions have unique physical characteristics that are permissive to initiating the double strand break (DSB) in the target genes. These characteristics may include the presence of non-B DNA forming sequences like the G-quadraplex structure, which becomes susceptible to substitutions in the DNA when the strand is no longer protected by torsional strain of the intertwined double helix^30,31,32^; as well as repetitive elements and palindromic sequences that may configure DSB susceptibility across the genome^33,34^, or through clustering of topological associated domains (TADs) within chromatin^35^. We have noted that CAD shares multiple sequence-specific binding sites at low frequency, which implies that structural elements may predispose the access/activity of CAD to the parent BPT gene bodies. Similarly, the capacity of CAD to generate DSBs versus single strand breaks in differentiating muscle cells also suggests an additional level of management over nuclease function.

Prior to our work here, studies have explored whether BPTs originate from activity of topoisomerases, enzymes that may induce DNA strand breaks through torsional strain, while converting DNA strands between relaxed and supercoiled states^36^. Early studies identified Topoisomerase I and II binding to oncogenic fusion genes that were causative for Burkitt’s lymphoma, mixed-lineage leukemia (MLL) and acute myeloid leukemia (AML)^37,38,39^. Recently, topoisomerase IIB has been linked to topologically susceptible BPTs that align with both chromosome loop structures and regions of paused RNA polymerase activity^40,41^. Nevertheless, a role of topoisomerases in DSB generation remains somewhat unclear, as these enzymes do not possess robust nuclease activity per se and may simply provide a relaxed DNA structure for DNase access.

One nuclease that has been identified as a functional prelude to BPT formation, is endonuclease G (EndoG). EndoG resides primarily within the mitochondria, however this nuclease is released from its organelle position and can traffic to the nucleus under cell stress where it fragments DNA, akin to the DNA damage associated with apoptosis^42^. Induction of replication stress has been shown to result in nuclear accumulation of EndoG, followed by DNA cleavage of the MLL BPT cluster^43^. EndoG has also been observed to act in concert with other DNA modifying enzymes, such as activation-induced cytidine deaminase (AID) to initiate MLL located DSBs, to propel the formation of MLL BPTs^44^. Whether EndoG initiates BPT formation on a larger genome scale remains unknown.

Together, our observations suggest that the initial step in the formation of an oncogenic BPT, the CAD-mediated generation of strand breaks in reciprocal gene pairs, is normal biology and is essential to lower the gene expression of factors, such as *Pax7*, that block the onset of a muscle cell differentiation program. We also demonstrate that CAD binds to several of the known BPT parent genes in the differentiating cell niche, BPTs that are known to cause leukemias, lymphomas and sarcomas. Finally, it is important to note that the data do not support the contention that BPT induced cancers solely originate in a self-renewing stem cell, an origin story referred to as the stem cell cancer hypothesis^45,46^. Rather our analyses suggest that the initial step in BPT formation begins with CAD targeting of susceptible loci in cells that have already initiated the differentiation program.

## Methods

### Animals

All mice were housed and treated at the University of Ottawa Animal Care and Veterinary Services in accordance with the Canadian Council on Animal Care (CCAC) guidelines and the University of Ottawa Animal Care Committee protocols.

### C2C12 Cell Culture Growth and Differentiation Protocols

C2C12 cells were obtained from the American Type Culture Collection (ATCC) and were initially provided in DMEM supplemented with 10% Fetal Bovine Serum (FBS) and 10% Dimethylsulphoxide (DMSO). The cells were thawed in a 37°C water bath and washed with 1X PBS to remove excess DMSO. Following this, the cells were centrifuged at 750 x g, resuspended in growth media (DMEM supplemented with 10% FBS and 2% Pen-Strep), and plated on 10 cm tissue culture plates. The cells were incubated at 37°C with 5% CO₂. Passaging was performed when cells reached 70% confluency, and the media was replaced every 48 hours to maintain cell growth. For differentiation, cells were induced under low serum conditions. The growth media was aspirated, cells were washed with 1X PBS, and differentiation media (DMEM supplemented with 2% Horse Serum and 2% Pen-Strep) was added. Cells were differentiated according to experimental protocols^3^.

### Primary Myoblast Isolation and Culture

Primary myoblasts were isolated from 4–6 week-old mice. For the isolation, skeletal muscle from the hind limbs was extracted. The exposed hind limbs were rinsed in 1X phosphate-buffered saline (PBS) and then cut into small pieces until a dispersed muscle isolate was obtained. The dispersed muscle lysate was cultured in a dispase/collagenase solution (Dispase II, Roche, and 1% collagenase reconstituted in sterile PBS with 2.5 mM CaCl₂), and to facilitate the enzymatic breakdown of the remaining tissue, multiple rounds of trituration were performed by pipette. This mixture was centrifuged at 750 x g for 5 minutes, and the pellet was resuspended in Ham’s F-10 Enriched Growth Media (Ham’s F-10 medium from Gibco Life Technologies, 20% FBS, 2% Penicillin-Streptomycin (Pen-Strep), and 2.5 μg/mL basicFGF (bFGF). This process yielded a population of primary myoblasts, at which point the cells were transferred to and grown on collagen-coated plastic tissue culture plates and maintained by replacing the Ham’s F-10 media every 48 hours to ensure optimal nutrient availability. Primary myoblasts were passaged at 70% confluency to prevent spontaneous differentiation. To induce differentiation, primary growth media was replaced with primary differentiation media consisting of Dulbecco’s Modified Eagle Medium (DMEM) supplemented with 2% horse serum and 2% Pen-Strep. Cells were harvested by scraping off the plates and centrifuged to pellet the cells at various growth and differentiation time points.

### Primary T Cell Isolation and Purification

T cells were isolated from the thymus of wild-type mice. The thymus was gently crushed in media and filtered through a single-cell filter. To purify the T cells, the EasySep Mouse Kit for positive selection was used, which selectively binds CD4 and CD8 cells to the magnet. The isolated cells were labeled with PE (phycoerythrin)-conjugated antibodies from the mouse. The remaining cells, double negative for CD4 and CD8, were collected and subjected to flow cytometry using the BD LSRFortessa™ Cell Analyzer with the following antibodies: T cell marker Thy1.2 (CD90), CD4, CD8, and a live/dead marker. Flow cytometry (FACS) confirmed that 70% to 80% of T cells were at the double-negative (DN) stage. The markers used for flow cytometry included CD4 (Anti-Mouse CD4, from Stemcell Technologies), CD8 (CD8 Monoclonal Antibody [53-6.7], from Invitrogen), and CD90.2 (a marker for thymus, BUV395 Rat Anti-Mouse from BD Biosciences) as described in common laboratory methods^47^.

### Plasmid Transfection and Generation of Stable Cell Lines

Plasmid transfections were performed as described^3^. Briefly, primary myoblasts were cultured until they reached 50-60% confluency in the absence of antibiotics. Purified plasmid DNA was prepared by diluting 2 μg of DNA and transfected using Lipofectamine 2000 (Invitrogen). The transfection mixture was incubated for 3 hours at 37°C. Following incubation, 1 mL of antibiotic-free growth media was added, and cells were allowed to grow overnight at 37°C. The next day, the growth media was replaced with fresh antibiotic-free growth media, which was changed every 48 hours post-transfection. For antibiotic selection, G418 (Invitrogen) was added to the growth media at a concentration of 400 μg/mL. Transfected cultures were maintained and passaged to generate stable pooled cultures.

### C2C12 Cell Culture Growth and Differentiation Protocols

C2C12 cells were obtained from the American Type Culture Collection (ATCC) and were initially provided in DMEM supplemented with 10% Fetal Bovine Serum (FBS) and 10% Dimethylsulphoxide (DMSO). The cells were thawed in a 37°C water bath and washed with 1X PBS to remove excess DMSO. Following this, the cells were centrifuged at 750 x g, resuspended in growth media (DMEM supplemented with 10% FBS and 2% Pen-Strep), and plated on 10 cm tissue culture plates. The cells were incubated at 37°C with 5% CO₂. Passaging was performed when cells reached 70% confluency, and the media was replaced every 48 hours to maintain cell growth. For differentiation, cells were induced under low serum conditions. The growth media was aspirated, cells were washed with 1X PBS, and differentiation media (DMEM supplemented with 2% Horse Serum and 2% Pen-Strep) was added. Cells were differentiated according to experimental protocols^3^.

### Caspase Inhibition Assays

For caspase-3 inhibition, cultured myoblasts were pre-treated with either 15 μM Z-DEVD-FMK (DEVD) from BioVision or DMSO for 4 hours at 37°C. After pre-treatment, cells were induced to differentiate using low serum media or continued in growth media, both supplemented with 15 μM DEVD or DMSO. The media was replaced every 48 hours throughout the differentiation period.

### ICAD Cloning and Site-Directed Mutagenesis

ICAD cloning constructs were previously described^3^. In brief, wild-type human ICAD was cloned into the reading frame A of the pcDNA3.1/HisC (Invitrogen) mammalian expression vector between the BamHI and NotI restriction sites. The PCR primers used included a BamHI site in the forward primer (5’-ATGGATCCGAGGTGACCGGGGACGCC-3’) a NotI site in the reverse primer (5’-TAGCGGCCGCCTATGTGGGATCCTGTCTG-3’). A two-step megaprimer approach was employed to introduce the D117E mutation. The forward primer listed above and a reverse primer (5’-CTGCCCCGCTCTCTGTTTCATC-3’) were used to generate the megaprimer, which was then purified and used with the original reverse primer to produce the full-length ICAD sequence containing the D117E mutation via PCR. The D224E mutation was introduced in a similar manner, using a reverse primer (5’-CTGATACCCGTCTCTACTGCATC-3’) to generate the megaprimer. The resulting PCR products were inserted between the BamHI and NotI restriction sites of pcDNA3.1/HisC as described above. All constructs were verified by sequencing.

### Fluorescent *in Situ* Hybridization (FISH)

Pax7 break-apart FISH probe kit (CT-PAC088, Cytotest), Vysis Foxo1 break-apart FISH probe kit (03N60-020, Abbott), and Pax7-Foxo1 fusion/translocation FISH probe kit (CT-PAC089, Cytotest) were used according to the manufacturer’s protocols (Fig.2d, Extended Data Fig. 2e). Briefly, primary myoblast cells were seeded on collagen-coated coverslips. Cells were fixed at both growth and 24 hours post-differentiation. The coverslips were then transferred to glass slides and immersed in denaturation solution (70% Formamide/2X SSC at 73 ± 1°C) for 5 minutes. The slides were dehydrated sequentially in 70% ethanol, 85% ethanol, and 100% ethanol for 1 minute each.

A probe mixture was prepared by combining 7 μL of hybridization buffer, 1 μL of probe, and 2 μL of purified water, which was vortexed, centrifuged, and incubated in a 73 ± 1°C water bath for 5 minutes. This mixture was applied to each coverslip to hybridize with the targeted DNA and incubated overnight (16 hours) at 37°C in a humidified chamber. After incubation, slides were washed with wash solution (2X SSC/0.1% NP-40) and air-dried in darkness before staining with DAPI (10 μL). For storage, slides were kept at 4°C and protected from light. For visualization, image z-stack acquisition was performed using a fluorescence microscope equipped with suitable filters. DAPI, SpectrumOrange/CytoOrange, and SpectrumGreen/CytoGreen fluorophores were used to visualize FISH signals. Images were then deconvolved using Zen Desk software.

### Protein Extraction and Western Blot

Primary myoblasts were isolated and proteins lysates were prepared as previously described^3^. Samples were loaded onto SDS-PAGE gels and then transferred to PVDF membranes. The membranes were blocked with 5% (w/v) non-fat powdered milk in TBS-T (10 mM Tris, pH 7.4; 150 mM NaCl; 0.05% Tween-20) and incubated overnight at 4°C with primary antibodies: rabbit anti-HIS tag (D3I10) mAb (12698, Cell Signaling); mouse anti-β-tubulin (clone E7, Developmental Studies Hybridoma Bank); and mouse anti-MF20 (Myosin Heavy Chain, Developmental Studies Hybridoma Bank). After incubation with primary antibodies, membranes were washed and incubated with HRP-conjugated secondary antibodies (goat anti-mouse or goat anti-rabbit, Bio-Rad). Protein expression was detected using an ECL detection kit (GE Healthcare). Relative protein levels were quantified using ImageJ software, with values normalized to α-tubulin as a loading control.

### Immunocytochemistry (ICC)

C2C12 cells were cultured on collagen-coated 25 mm coverslips in 6-well Petri plates. At designated time points, the culture media was removed by aspiration and cells were washed three times with 1X PBS. Cells were fixed with 4% paraformaldehyde (PFA) in PBS for 10 minutes, then quickly washed twice with 1X PBS. Fixed cells were incubated with a blocking solution of 5% bovine serum albumin (BSA) in 1X PBS for 1 hour, followed by overnight incubation with the primary antibody at 4°C. The next day, cells were washed and incubated with a fluorescently labeled species-specific secondary antibody for the corresponding primary antibody, as indicated in the figure legends. The secondary antibody was applied for 1 hour in the dark. Cells were then washed and counterstained with DAPI (1:10,000 in 1X PBS) for 10 minutes to stain the nuclei. The labeled coverslips were mounted onto glass microscope slides using a drop of Dako Fluorescent Mounting Medium (Dako). Cells were visualized using a Zeiss Observer Z1 Inverted Microscope with appropriate filter settings.

### RNA Isolation and Quantitative Reverse Transcription, Real Time PCR

RNA was isolated from primary myoblast cells during growth conditions, and for the indicated times post-low serum induction of differentiation, using the RNeasy Mini Kit (Qiagen). A total of 1 μg RNA was then used to synthesize cDNA through reverse transcription with the iScript cDNA synthesis Kit (BioRad). Quantitative gene expression analysis of Pax7 and the Pax7-Foxo1 fusion was performed using quantitative RT-PCR (qPCR). The cDNA was subjected to qPCR using SYBR® Green Supermix (BioRad) with 5 μM forward and reverse primers in an Eco™ Real-Time PCR system (Illumina). qPCR primers were synthesized as follow: *Pax7*-forward 5’-GTT TAT CCA GCC GAC TCT GG-3’; *Pax7*-reverse 5’-ATC GGC ACA GAA TCT TGG AG-3’; *Foxo1*-forward 5’-TAT TGA GCG CTT GGA CTG TG-3’; *Foxo1*-reverse 5’-TGG ACT GCT CCT CAG TTC CT-3’; *GAPDH*-forward 5’-TTG GGT TGT ACA TCC AAG CA-3’; and *GAPDH*-reverse 5’-CAA GAA ACA GGG GAG CTG AG-3’). For the amplification of the *Pax7-Foxo1* fusion transcript, *Pax7* forward primer and *Foxo1* reverse primer were used for the PCR. Gene expression analysis was performed using a two-step PCR cycle. The run began with a 5-minute hot start activation at 95°C, followed by 40 cycles of 10-second denaturation at 95°C and a 30-second combined annealing/extension step at 60°C. The critical threshold (CT) was set to encompass the exponential amplification curve of each sample. The delta CT (ΔCT) was calculated by subtracting the CT of the gene of interest from that of the GAPDH control, and values were normalized as indicated in the corresponding legends.

### Statistical analysis

Statistical quantification was performed using Prism (version 10.0.3). All values are expressed as arithmetic mean ± SEM. Statistical significance in (Extended Data Fig.2d) was determined using one-way ANOVA, with Dunnett’s T-test and in (Fig.2f) was determined using two-way ANOVA, with Tukey’s multiple comparison test. For all experiments, *: p<0.05, **: p<0.01, ***: p<0.001****: p<0.0001.

### RNA-sequencing

Total RNA was isolated from primary mouse myoblasts (age of 8-12 weeks). Cells were harvested by scraping off the plates and centrifuged to pellet the cells at growth and (differentiation time points 12,24,48, and 72). For RNA isolation, the NucleoSpin RNA kit from Macherey-Nagel (cat. 740990.50) was used. Libraries were prepared and sent to the IRCM sequencing facility. Sequencing was performed on an Illumina NovaSeq 6000.

### Cleavage Under Targets and Tagmentation (CUT&Tag)

CUT&Tag experiments were performed following the steps previously described by Li et al.^48^ as adapted from the original protocol^49^. Briefly, fresh C2C12 cells or primary T cells were immobilized on Concanavalin A-coated magnetic beads and permeabilized with digitonin. Cells were incubated with primary antibody against CAD (anti-Dff40 rabbit Ab, AB16926, Millipore) or with rabbit IgG from Santa Cruz (SC-2027). Tethered cells were washed thoroughly to limit unbound antibodies, then incubated with pA-Tn5 fusion protein for an hour. Tn5 transposase with the addition of MgCl2 induced tagmentation of chromatin. Then tagmented DNA was extracted and amplified by PCR to prepare libraries for sequencing on Illumina NovaSeq 6000.

### Bioinformatics Analysis for CUT&Tag and RNA-Seq Data Processing from Next-Generation Genome Sequencing

First, quality control (QC) of raw fastq.gz reads was observed using FastQC while QC reports of processed reads in subsequent steps were compiled with MultiQC. Cutadapt was used to trim Illumina adapter sequences from the raw reads. Bowtie2 was used to map the trimmed reads to the reference genome and generate the alignment (.bam) files for CUT&Tag while Hisat2 was used for RNASeq. Picard and SAMtools sorted the alignments by coordinate, marked duplicates, and filtered out low-quality reads below Q value of 10. The final filtered alignment was indexed using SAMtools and contains only mapped reads without duplicates for CUT&Tag, while retaining duplicates for RNASeq. Coverage was computed using deepTools bamCoverage with highest resolution (binSize of 1) and normalized by RPGC (and BPM) to obtain bigwig and bedgraph files. Peak calling of CUT&Tag binding sites was performed by MACS, SEACR, and GoPeaks. Heatmap profiles were created using deepTools computeMatrix with “scale-regions” and “skipZeros” options followed by plotHeatmap and plotProfile with “perGroup” option. The accession numbers for associated data are: CUT and Tag is GSE277954, and RNA-Seq. is GSE277956.

### Data Processing for Differential Binding Analysis

DiffBind was used to perform pair-wise differential (and shared) binding analysis. An “occupancy analysis” was performed using the peak files for each condition. Consensus peak sets were produced for each condition in the pair-wise comparison. Peaks which overlap in 2/3 peak callers were retained in a caller consensus set for each replicate. Of these, peaks which overlap in at least 2 replicates from the caller consensus sets were retained as a final high-confidence consensus peak set for that condition. A differential comparison of these final consensus peak sets between conditions were made. Finally, these differential and shared peaks were annotated with ChIPseeker using the UCSC database of known genes. Additionally, KEGG and GO analysis were performed using clusterProfiler from the resulting gene annotations.

### Data Processing for Circa and UpSet Plots

Firstly, a CAD gene set was obtained by removing IgG consensus peaks and ENCODE blacklisted regions from the CAD-DEVD-24 MACS/SEACR/GoPeaks consensus peak set, followed by annotation of the resulting peak set. Secondly, the set of fusion genes from the COSMIC (https://cancer.sanger.ac.uk/cosmic) database were mapped to their mm10 orthologs using bioMART. Finally, the intersection and union of these CAD and COSMIC gene sets were plotted using the UpSetR package’s *upset* function.

Circa plots were made with the circlize package (R package). The “CAD-Fusion genes” were obtained from the union of CAD and COSMIC gene sets determined above and were plotted as the outer ring in the circa plot. Breakpoints from the COSMIC database were mapped to the mm10 genome assembly using the liftOver tool (in R). These breakpoints were filtered to retain only those within the ranges of the CAD-Fusion gene regions. CAD peaks overlapping these breakpoints in the CAD-Fusion gene set were determined using ChIPseeker’s *findOverlapsOfPeaks* function. These CAD sites overlapping COSMIC breakpoints were then plotted along the inner ring of the circa plot. Any fusion gene pairs where both genes were within the CAD-Fusion gene set were also plotted with ribbon connections in the circa plot.

### Data Processing for Motif Analysis

Sequences for the peak regions of IgG-filtered CAD sites were obtained from the UCSC mm10 hard-masked assembly. These CAD site sequences were passed through STREME for motif discovery^50^. Subsets of the CAD sites (i.e. those found in promoter regions, COSMIC genes, and overlapping breakpoints) were also analyzed against the remaining sites as a control. Additionally, a separate motif analysis was performed on the same CAD sites in which the “sequence preference” was determined by the nucleotide frequency across the regions. This method and main code was adopted from Gnugge et al.^51^.

### Data processing for RNAseq

Raw gene counts were obtained from the alignment files using *featureCounts*. Differential expression analysis and statistics were obtained from DESeq2 results by performing pairwise comparisons of each differentiation timepoint with the growth timepoint as reference. Genes with p.adjusted < 0.05 across all timepoint comparisons were kept for the expression trend analysis. An expression trend was then determined if the log2FoldChange of subsequent sequential timepoints was continuous (greater for increasing expression trend, lower for decreasing expression trend) for each gene. Additional details for the replicates numbers and conditions are explained in the corresponding legend in the figure.

## Acknowledgements

The authors would like to thank members of the Megeney and Dilworth laboratories and Sandra Megeney for helpful discussions. We would also like to thank Hina Bandukwala for assistance with the establishment of the bioinformatic analyses pipelines and Sarah Boissel for CUT&Tag library sequencing. The work in this publication was supported by grants to L.A.M. from the Canadian Institutes of Health Research (CIHR) (PJT-156120 and PJT-175274), to F.J.D. and L.A.M. from CIHR (TGH-158224), to M.B. from the National Institutes of Health (2R01DK098449-06) and by a graduate scholarship to D.A. from King Saud University, Saudi Arabia.

---

**Extended Data Figure 1.**
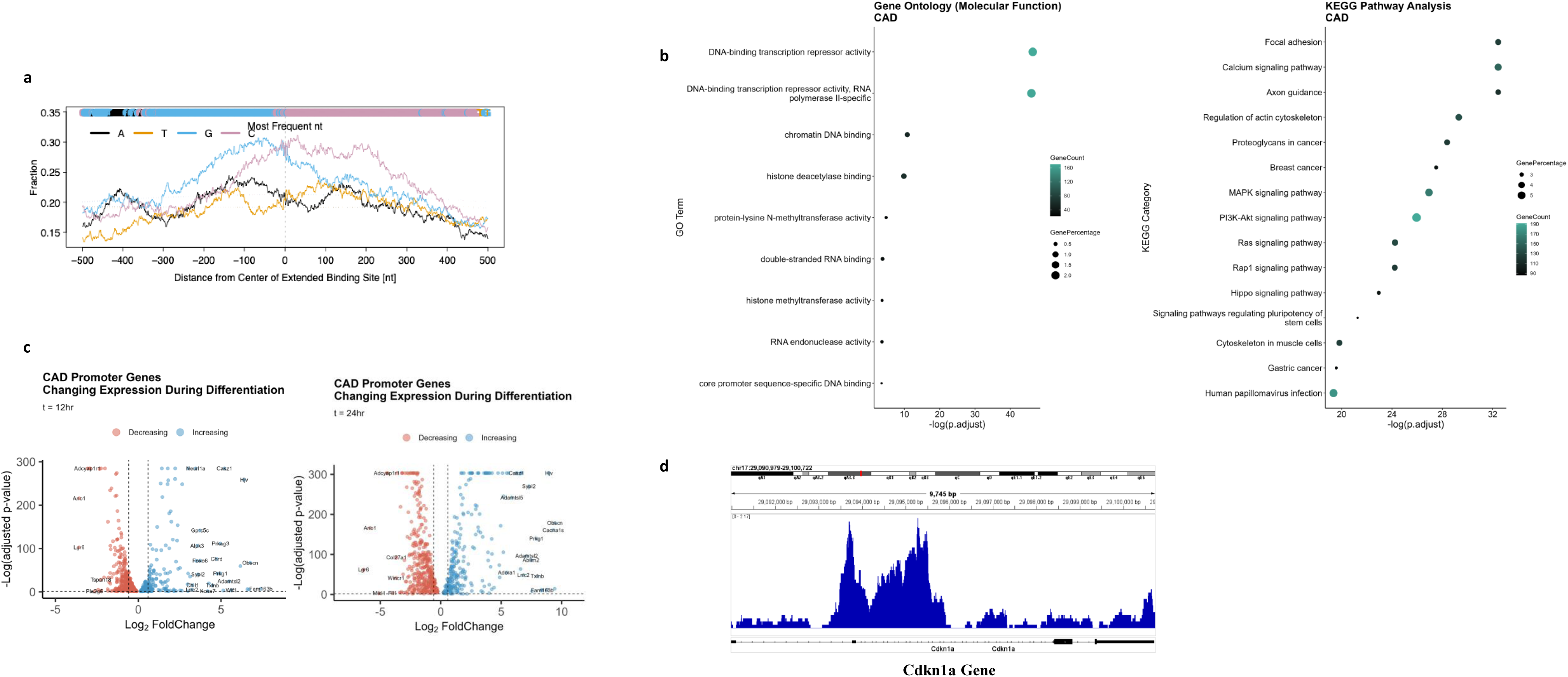
Gene expression analysis of CAD bound genes in differentiating muscle cells. (A) All peaks for CAD are called by MACS regions extended +/− 500bp from their summits. Followed by mapping them to masked mm10 genome to get nucleotide sequences. The alignments have higher representation of GC-rich sequences in CAD binding genome. (B) KEGG and Gene Ontology (GO) analysis identified within CAD functional genes Top 15 terms are enriched for molecular functions. (C) The volcano plot shows the genes with trending expression changes at 12hr and 24 hr differentiation for which CAD is bound to those gene promoters. (D) Integrative genomics viewer (IGV) snapshots depicting CAD CUT&Tag signal associated peak calls for the Cdkn1a gene from C2C12 skeletal muscle cells.

**Extended Data Figure 2.**
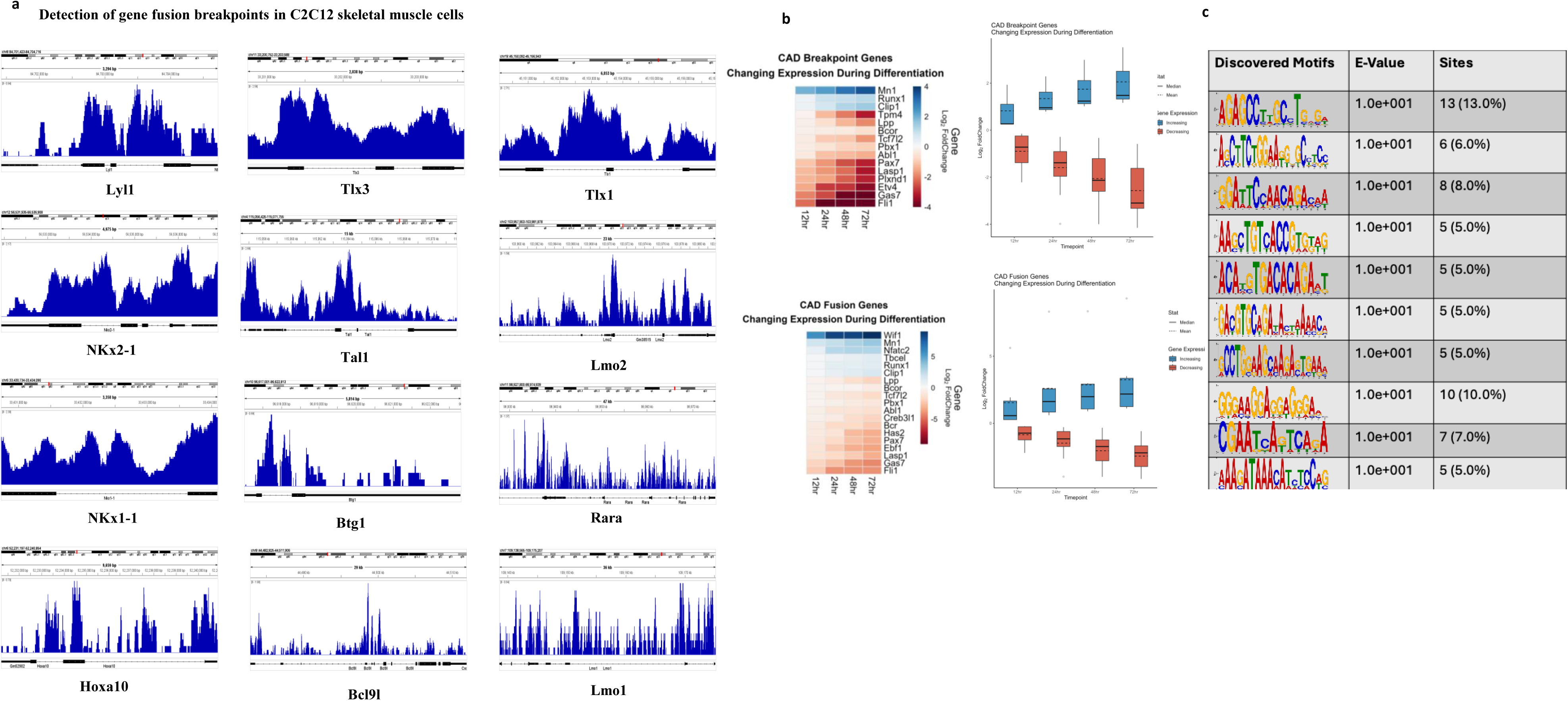
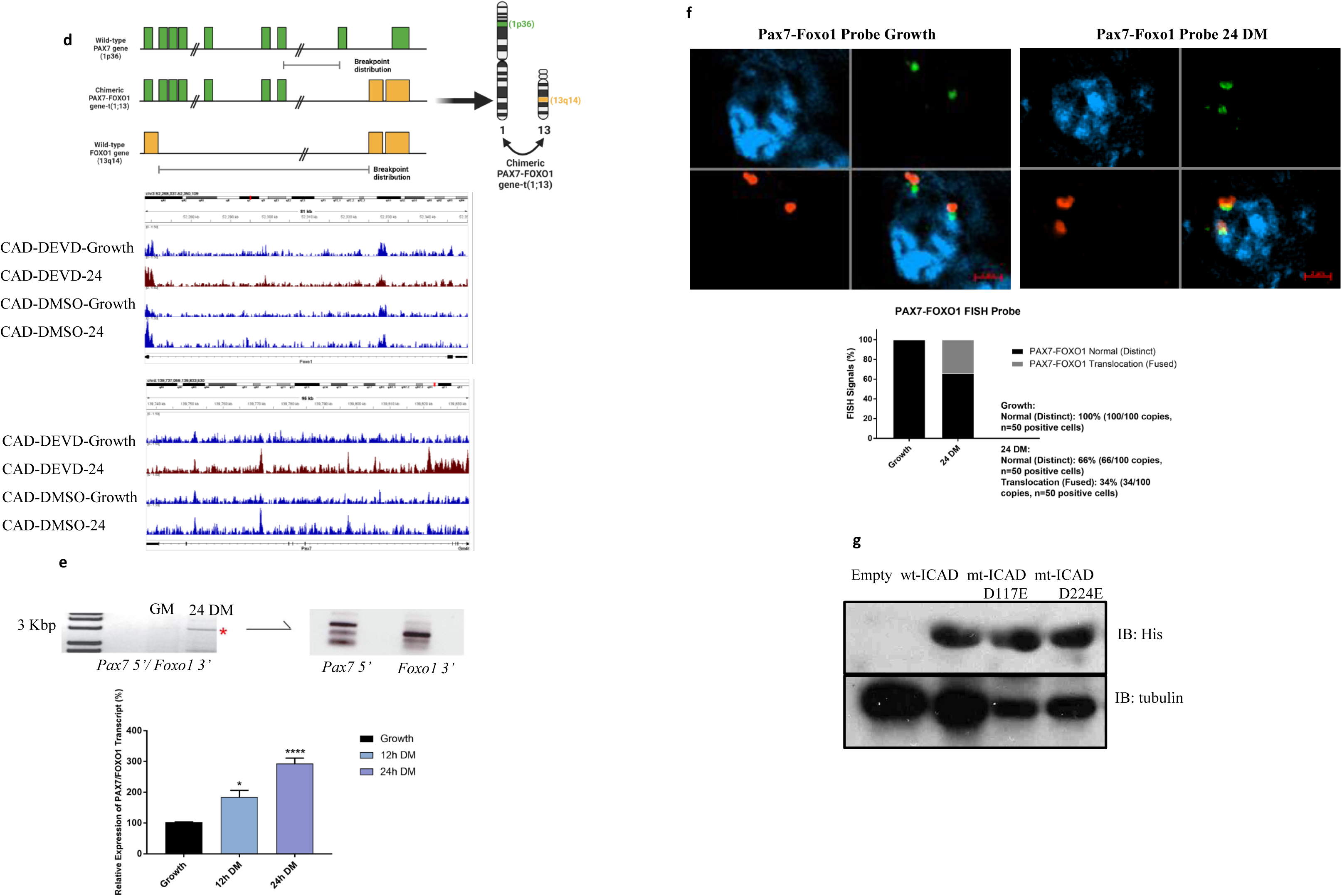
CAD targeted breakpoint translocation (BPT) genes in differentiating C2C12 muscle cells. (A) IGV snapshots depicting CAD CUT&Tag signal and associated peak calls for indicated genes in C2C12 skeletal muscle cells. (B) Heatmaps depicting changes in gene expression levels in CAD fusion genes or CAD breakpoint genes during 12-, 24-, 48-, and 72-hours differentiation. Blue color indicates elevated mRNA levels, while red color indicates repressed levels. Color scale represents the log2fold change. Three biological replicates for each timepoint were assessed. Boxplot representing CAD breakpoint genes or CAD fusion genes changing expression during differentiation. (C) Graphical representation of sequence motifs detected in the CAD bound genome using Motif enrichment analysis calculated with STREME; E-values of motifs are indicated). (D) Schematic outline depicting the chromosomal translocations producing the chimeric proteins PAX7/FOXO1 t(1;13). Integrative genome view representing enrichment of CAD at Pax7 and Foxo1 genomic loci. Genome tracks identified by MACS2 peak caller of CAD treated with DMSO or DEVD (growth and 24 hours post-differentiation). (E) Detection of Pax7-Foxo1 fusion transcript in differentiating primary myoblasts. Primers specific to the 5’ region of the Pax7 transcript and the 3’ region of the Foxo1 transcript were used to probe cDNA libraries of differentiating primary myoblasts for the appearance of a fusion transcript. PCR product was observed at ∼3000 bp that corresponded to the expected size of a Pax7-Foxo1 transcript. Secondary PCR from the isolated 3000bp band was performed for the 5’ region of the Pax7 transcript and 3’ region of the Foxo1 transcript. (N=2). Quantitative PCR (N=4) of Pax7-Foxo1 gene fusion during 12- and 24 hours post-differentiation. *: p<0.05, ****: p<0.0001 when compared to growth using one-way ANOVA with Dunnett’s multiple comparisons test. (F) FISH analysis detect Pax7-Foxo1 gene fusions in primary myoblast cells during cell growth and 24 hours post differentiation using PAX7-FOXO1 Fusion/Translocation FISH Probe Kit (CT-PAC089, Cytotest). The PAX7-FOXO1 fusion FISH probe set is designed to detect rearrangements between the human *PAX7* and *FOXO1* genes, located on chromosome band 1p36.13 and 13q14.11, respectively. For Pax7-Foxo1 FISH signals, 100 copies (n=50) of distinct FISH probes (100%) were quantified during growth condition. In differentiated conditions (n=50), 66 copies of distinct FISH probes (66%) and 34 copies of fused (translocated) FISH signals (34%) were quantified. (G) Western blot analysis of His-tagged ICAD constructs were detected with a mouse anti-His primary antibody in transfected mouse primary myoblasts (Tubulin was used to control for loading).

**Extended Data Figure 3.**
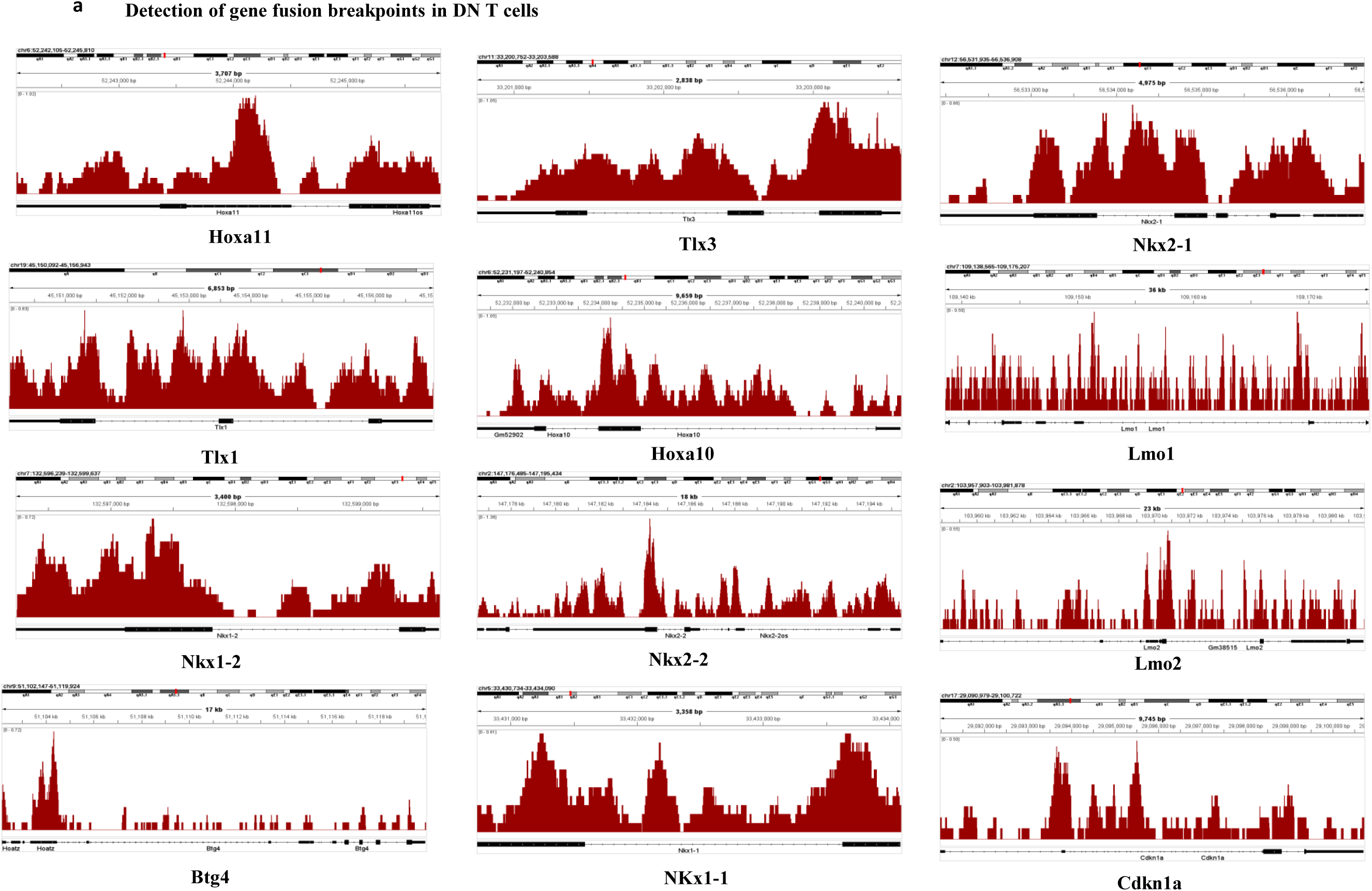
CAD targeted breakpoint translocation (BPT) genes in early differentiating T cell progenitors. (A) IGV snapshots depicting CAD CUT&Tag signals and associated peak calls for breakpoint translocation (BPT) genes identified in early differentiating T cell progenitors.

